# Calcium influx rapidly establishes distinct spatial recruitments of Annexins to cell wounds

**DOI:** 10.1101/2023.12.03.569799

**Authors:** Mitsutoshi Nakamura, Susan M. Parkhurst

**Affiliations:** Basic Sciences Division, Fred Hutchinson Cancer Center, Seattle, WA, USA 98109

**Keywords:** cell wound repair, Drosophila, Annexin, calcium dynamics

## Abstract

To survive daily damage, the formation of actomyosin ring at the wound periphery is required to rapidly close cell wounds. Calcium influx is one of the start signals for these cell wound repair events. Here, we find that rapid recruitment of all three *Drosophila* calcium responding and phospholipid binding Annexin proteins (AnxB9, AnxB10, AnxB11) to distinct regions around the wound are regulated by the quantity of calcium influx rather than their binding to specific phospholipids. The distinct recruitment patterns of these Annexins regulate the subsequent recruitment of RhoGEF2 and RhoGEF3 through actin stabilization to form a robust actomyosin ring. Surprisingly, we find that reduced extracellular calcium and depletion of intracellular calcium affect cell wound repair differently, despite these two conditions exhibiting similar GCaMP signals. Thus, our results suggest that, in addition to initiating repair events, both the quantity and sources of calcium influx are important for precise Annexin spatiotemporal protein recruitment to cell wounds and efficient wound repair.

**Summary:** Cells have rapid and robust repair systems to survive daily damage. This study shows that calcium influx regulates the three distinct *Drosophila* Annexin recruitment patterns to the cell wound in order to organize an actomyosin ring for efficient wound closure.

## INTRODUCTION

When cells encounter physical stresses that result in cortex ruptures, they must rapidly deploy their repair pathway to preserve cell homeostasis. Influx of extracellular calcium through the cortex breach is one of the most upstream events and is thought to initiate the repair process by activating downstream molecules (Andrews and Corrotte, 2018; Cheng et al., 2015; Cooper and McNeil, 2015; Ebstrup et al., 2021; Moe et al., 2015; Nakamura et al., 2018). Previous studies have shown that calcium influx as detected by calcium indicators (i.e., GCaMP) is uniform at the wound site (Davenport et al., 2016; Nakamura et al., 2020; Terasaki et al., 1997). How this uniform calcium signal is translated into the dynamic, yet precise, spatial and temporal recruitment and activity of downstream molecules is unknown. Upon cortex injury, vesicles are recruited to the wound forming a temporary membrane plug that seals the damaged area, followed by the assembly of an actomyosin ring that links the cortical cytoskeleton to the plasma membrane at the wound edge which then translocates inward to pull the wound edges closed (Cooper and McNeil, 2015; Hui et al., 2022; Nakamura et al., 2018; Sonnemann and Bement, 2011). After the wound closes, the plug is removed from the wound site, the actomyosin ring is disassembled, and the cortical cytoskeleton and plasma membrane are remodeled to return their original composition, organization and functional state.

Annexins (Anxs) are a highly conserved family of calcium responding proteins that bind specific phospholipids calcium-dependently (Blackwood and Ernst, 1990; Gerke et al., 2005; Lizarbe et al., 2013). Anxs are known to play roles in wound repair: AnxA1, AnxA2, AnxA4, AnxA5, AnxA6, and AnxA7 (out of 13 mammalian Anxs) are rapidly recruited (within seconds) to wounds and associate with the plasma membrane to regulate different aspects of cell wound repair context-dependently (Bittel et al., 2020; Carmeille et al., 2016; Croissant et al., 2020; Davenport et al., 2016; Koerdt and Gerke, 2017; Lennon et al., 2003; McNeil et al., 2006; Nakamura et al., 2017; Pervin et al., 2018; Sønder et al., 2019; Vicic et al., 2022). In MCF7 cells, AnxA4 has been suggested to induce membrane curvature and AnxA6 to induce membrane folding, which are necessary to generate the constriction force needed to pull the wound edge inward to close the gap (Boye et al., 2017). AnxA5 has been shown to form a 2D self-assembled lattice structure at wounds to reseal the damaged plasma membrane in mouse perivascular cells (Bouter et al., 2011). AnxA1, AnxA2, AnxA5, and AnxA6 are recruited to muscle cell wounds where they assemble into a compact structure on the extracellular surface of a cell, called a tight repair cap, for efficient wound repair (Demonbreun et al., 2016). AnxA6 has been shown to form a similar tight repair cap for efficient membrane repair in neurons (Demonbreun et al., 2022). In addition to dynamic membrane regulation, AnxA1, AnxA2, and AnxA6 have also been shown to regulate actin dynamics through binding to and/or bundling F-actin (Glenney et al., 1987; Hayes et al., 2004; Hosoya et al., 1992; Ikebuchi and Waisman, 1990). In particular, AnxA2 binds to F-actin and regulates Rho-mediated actin rearrangement during cell adhesion and cytokinesis, and regulates actin polymerization and bundling during exocytosis and endosome biogenesis (Babbin et al., 2007; Benaud et al., 2015; Benaud and Prigent, 2016; Gabel et al., 2015; Morel et al., 2009; Rescher et al., 2008).

Using the *Drosophila* cell wound repair model, we have previously shown that AnxB9 is rapidly recruited to the wound where it regulates actin stabilization, which is necessary for the recruitment of RhoGEF2 to wounds during the repair process (Nakamura et al., 2017). Here we show that the other two *Drosophila* Anxs, AnxB10 and AnxB11, are also recruited to wounds, where they are both required for RhoGEF3 recruitment to wounds. Strikingly, all three Anxs are recruited to wounds within 3 seconds post-wounding, exhibit distinct but adjacent spatial recruitment patterns, and have non-redundant functions in actomyosin ring formation during cell wound repair. Interestingly, while Anxs are not recruited to wounds in the absence of a calcium influx, we find that reduced calcium influx alters their recruitment patterns. Thus, our results indicate that calcium influx upon wounding not only triggers the repair process but creates the distinct spatiotemporal protein recruitment patterns of molecules such as the Anxs.

## RESULTS AND DISCUSSION

### All three *Drosophila* Annexins are rapidly recruited to wounds and exhibit distinct spatial recruitment patterns

We previously showed that the rapid spatial and temporal recruitment of RhoGEF2 to wounds is regulated by AnxB9 (Fig. 1A-B’’) (Nakamura et al., 2017). Since RhoGEF3 and Pbl are still recruited to wounds in the AnxB9 knockdown background, we investigated if the other two *Drosophila* Anxs—AnxB10 and AnxB11—regulate the spatial and/or temporal recruitment of these two RhoGEFs during cell wound repair. If so, both AnxB10 and AnxB11 would be expected to be recruited to wounds before RhoGEF patterns are established (<15 sec post wounding). To examine the response of AnxB10 and AnxB11 during cell wound repair, we generated three fly transgenic lines expressing: (1) sfGFP-AnxB10 driven by the Myosin regulatory light chain (Spaghetti Squash (Sqh)) promoter, (2) mScarlet (StFP)-AnxB10 (knock-in), and (3) StFP-AnxB11 under the control of the endogenous AnxB11 promoter. We examined embryos co-expressing these fluorescently-tagged Anxs and an actin reporter (sGMCA or sStMCA; see Methods). Wounds were generated by laser ablation on the lateral side of nuclear cycle 4-6 *Drosophila* embryos (see Methods). In this cell wound repair model, actin accumulates in two adjacent regions: 1) a highly-enriched ring at the wound edge, and 2) a less dense halo region encircling the actomyosin ring at the wound periphery (Fig. 1A) (Abreu-Blanco et al., 2011).

**Figure. 1.**
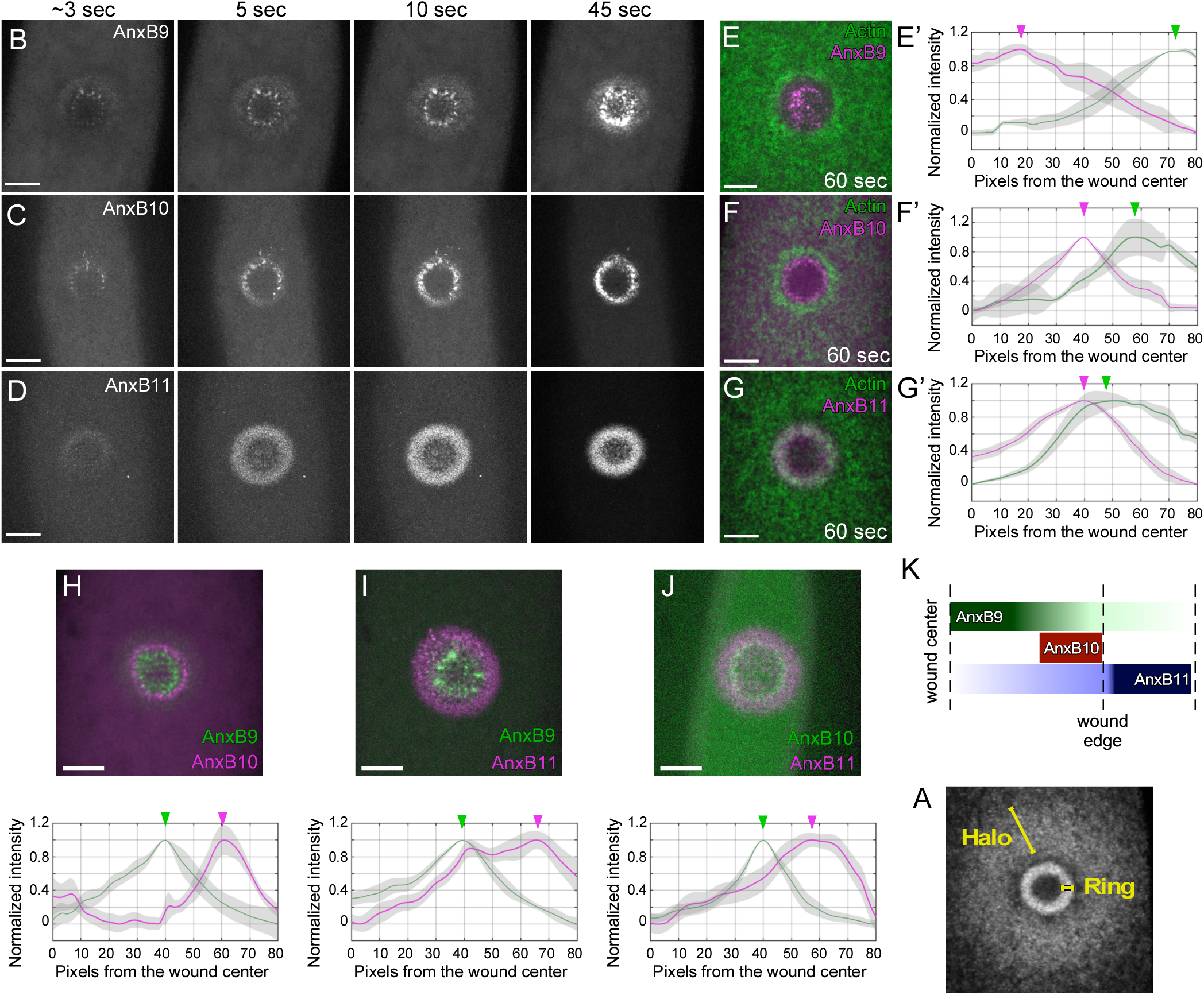
The three *Drosophila* Anxs are rapidly recruited to distinct regions around the wound. (A) Confocal xy projection image from a wounded NC4-6 *Drosophila* embryo expressing an actin reporter (sGMCA). (B-D) Confocal xy projection image from a wounded NC4-6 *Drosophila* embryo expressing GFP-AnxB9 (B), sfGFP-AnxB10 (C), or StFP-AnxB11 (D). (E-G) Confocal xy projection image from a wounded NC4-6 *Drosophila* embryo expressing an actin reporter (sGMCA or sStMCA) and GFP-AnxB9 (E), StFP-AnxB10 (F), or StFP-AnxB11 (G). (E’-G’) Normalized fluorescence intensity profile at 60 sec post wounding from ≥10 embryos expressing sStMCA/GFP-AnxB9 (E’; n=13), sGMCA/AnxB10 (F’; n=10), and sGMCA/StFP-AnxB11 (G’; n=10). Fluorescence peaks are indicated by arrowheads. (H-J) Confocal xy projection image from a wounded NC4-6 *Drosophila* embryo expressing GFP-AnxB9/StFP-AnxB10 (H), GFP-AnxB9/StFP-AnxB11 (I), and sfGFP-AnxB10/StFP-AnxB11 (J). (E’-G’) Normalized fluorescence intensity profile at 60 sec post wounding from ≥10 embryos expressing GFP-AnxB9/StFP-AnxB10 (H’; n=10), GFP-AnxB9/StFP-AnxB11 (I’; n=11), and sfGFP-AnxB10/StFP-AnxB11 (J’; n=13). (K) Schematic diagram summarizing the localization patterns of three Anxs at the wound edge. Scale bar: 20 µm. Time after wounding is indicated. In fluorescence intensity profiles, the line represents the averaged fluorescent intensity, and the gray area is the 95% confidence interval. Fluorescence peaks are indicated by arrowheads.

Upon wounding, both AnxB10 and AnxB11 are rapidly recruited to wounds (<3 sec), similar to that observed with AnxB9 (Fig. 1B-D) (Nakamura et al., 2017). While their temporal recruitment is similar, the three Anx spatial recruitments to wounds differ. AnxB9 recruitment forms two distinct ring-like structures: a thin discontinuous ring with punctate accumulation inside the actin ring and a wide diffuse outer ring whose inner edge overlaps with the actin ring (Fig. 1B, E-E’). AnxB10 is recruited in a sharp ring that is inside and juxtaposed to the actin ring (Fig. 1C, F-F’). AnxB11 is recruited in a robust ring that overlaps with the actin ring and is also recruited to the area inside of the actin ring in a diffuse distribution (Fig. 1D, G-G’). These distinct Anx spatial recruitment patterns were confirmed when examining different combinations of fluorescently-tagged Anxs (Fig. 1H-J’). Overall, the three Anxs exhibit distinct spatial recruitment patterns compared to each other (Fig. 1K), but which are similar to those observed for the three RhoGEFs that they regulate (Nakamura et al., 2017).

### *Drosophila* Annexins have non-redundant functions in cell wound repair

The distinct spatial recruitment patterns of Anxs to wounds suggest that each Anx has a different role during cell wound repair. To investigate this, we examined wound repair dynamics of embryos expressing an actin reporter (sGMCA; (Kiehart et al., 2000)) in each Anx RNAi knockdown background. As previously described, AnxB9 knockdown embryos exhibit wound overexpansion, slow wound repair, decreased actin ring width, and lower actin accumulation in the actin ring (Fig. 2A-B’, G, L-O; Table S2; Video 1) (Nakamura et al., 2017). AnxB10 knockdown embryos also exhibit wound overexpansion, slow wound repair, decreased actin ring width, and lower actin accumulation in the actin ring (Fig. 2C-C’, H, L-P; Fig. S1A-A’; Table S2; Video 1). While AnxB11 knockdown embryos exhibit wound overexpansion and lower actin accumulation in the actin ring, contraction rates and actin ring width are similar to control embryos (Fig. 2D-D’, I, L-P; Fig. S1B-B’; Table S2; Video 1). Thus, the recruitment patterns of Anxs to wounds and their knockdown phenotypes indicate that each Anx acts non-redundantly in regulating actin ring dynamics during cell wound repair.

**Fig. 2.**
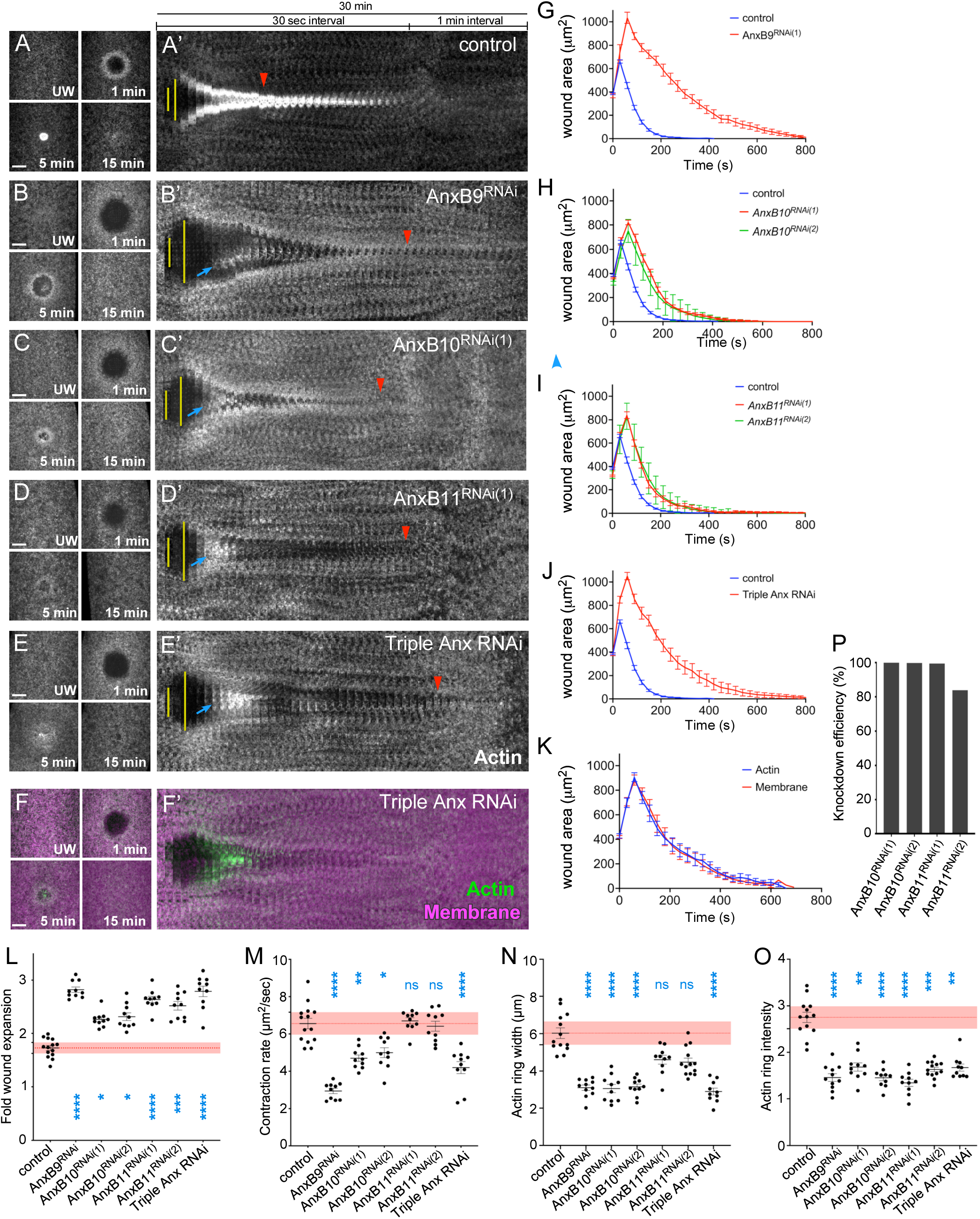
The three Drosophila Anxs have non-redundant functions during cell wound repair. (A-E) Actin dynamics (sGMCA) during cell wound repair in NC4–6 staged *Drosophila* embryos: control (A), AnxB9 RNAi (B), AnxB10^RNAi(1)^ (C), AnxB11^RNAi(1)^ (D), triple Anx RNAi (E). Wound expansion is highlighted by yellow lines. Actin accumulation internal to the wound is indicated by blue arrows. Completion of wound closure is indicated by red arrowheads. (F) Actin and membrane dynamics (sGMCA and StFP-8PM, respectively) during cell wound repair in triple Anx knockdown embryos. (G-J) Quantification of the wound area over time for control (n=14), AnxB9 RNAi (G; n=10), AnxB10^RNAi(1)^ (H; n=10), AnxB11^RNAi(1)^ (I; n=10), triple Anx RNAi (J; n=10). (K) Quantification of the wound area over time measured from actin (sGMCA) and membrane (StFP-8PM) reporters in triple Anx knockdown embryos (n=10). (L-O) Quantification of fold wound expansion (L), contraction rate (M), actin ring width (N), and actin ring intensity (O) for control, AnxB9 RNAi, AnxB10 RNAi, AnxB11 RNAi, triple Anx RNAi. (P) qPCR analysis of knockdown efficiency in AnxB10^RNAi(1)^, AnxB10^RNAi(2)^, AnxB11^RNAi(1)^, and AnxB11^RNAi(2)^ embryos under our imaging conditions. Error bars represent ± SEM. Scale bar: 20 µm. Time after wounding is indicated. ANOVA tests were performed in L-O: * *p*<0.05, ** *p*<0.01, *** *p*<0.001, **** *p*<0.0001, ns is not significant.

Since a previous study in mammalian tissue culture cells suggested that Anxs constrict the plasma membrane breach through binding directly to the plasma membrane during cell wound repair (Boye et al., 2017; Davenport et al., 2016), we next examined whether *Drosophila* Anxs could close plasma membrane wounds independently of actin ring closure. To visualize the plasma membrane in our system, we wounded embryos co-expressing a plasma membrane reporter (Ubi-StFP-8PM), along with an actin reporter, in a triple Anx knockdown (AnxB9 RNAi + AnxB10 RNAi + AnxB11 RNAi) background. We find that closure of both the actin ring and plasma membrane breaches are still highly correlated in the triple Anx knockdown background, with these triple knockdown embryos exhibiting wound overexpansion, slow wound repair, decreased actin ring width, and lower actin accumulation in the actin ring (Fig. 2E-F’, J-O; Table S2; Video 1). Hence, in the context of *Drosophila* cell wound repair, it is unlikely that Anxs regulate membrane dynamics actin-independently to close wounds.

### AnxB10 and AnxB11 regulate RhoGEF2 and RhoGEF3 recruitment to cell wounds

Since AnxB10 and AnxB11 are recruited to wounds earlier than RhoGEFs, have similar spatial recruitment patterns to wounds as RhoGEFs, and have non-redundant functions for actin ring dynamics during cell wound repair, we expected that AnxB10 and AnxB11 would regulate RhoGEF3 and Pbl recruitment to wounds, respectively. We wounded embryos co-expressing sfGFP-RhoGEF3 or Pbl-sfGFP, along with an actin reporter, in AnxB10 or AnxB11 knockdown backgrounds (Fig. 3A-H). In control embryos, RhoGEF3 is recruited to the actin ring and outside the ring, and Pbl is recruited outside the actin ring (Fig. 3A, F). Surprisingly, Pbl is still recruited to wounds in both AnxB10 and AnxB11 knockdown backgrounds, indicating that its regulation does not depend on Anxs. In contrast, RhoGEF3 is no longer recruited to wounds in either of the AnxB10 and AnxB11 knockdown backgrounds (Fig. 3A-C, F-H, N-O). To determine if AnxB10 and AnxB11 regulate RhoGEF3 recruitment to the wound by stabilizing actin as AnxB9 does for RhoGEF2, we examined RhoGEF3 recruitment to the wound in embryos injected with phalloidin (actin stabilizer). Intriguingly, the recruitment of RhoGEF3 to the wound in AnxB10 and AnxB11 knockdown backgrounds is partially rescued by phalloidin injection, suggesting that both of these Anxs work upstream of RhoGEF3 through actin stabilization (Fig. 3D-E, N). Since RhoGEF2 recruitment to the wound also depends on actin stabilization, we wounded embryos co-expressing sfGFP-RhoGEF2 and the actin reporter in AnxB10 and AnxB11 RNAi backgrounds. As expected, RhoGEF2 is no longer recruited to the wound in AnxB10 and AnxB11 RNAi backgrounds (Fig. 3I-K, P). Surprisingly, in this context RhoGEF2 recruitment is not rescued by phalloidin injection (Fig. 3L-M, P). Thus, AnxB10 and AnxB11 must regulate another aspect of actin dynamics in addition to actin stabilization during cell wound repair. Taken together, our results indicate that the three Anxs regulate spatial recruitment of RhoGEF2 and RhoGEF3 to wounds during cell wound repair through controlling actin dynamics.

**Fig. 3.**
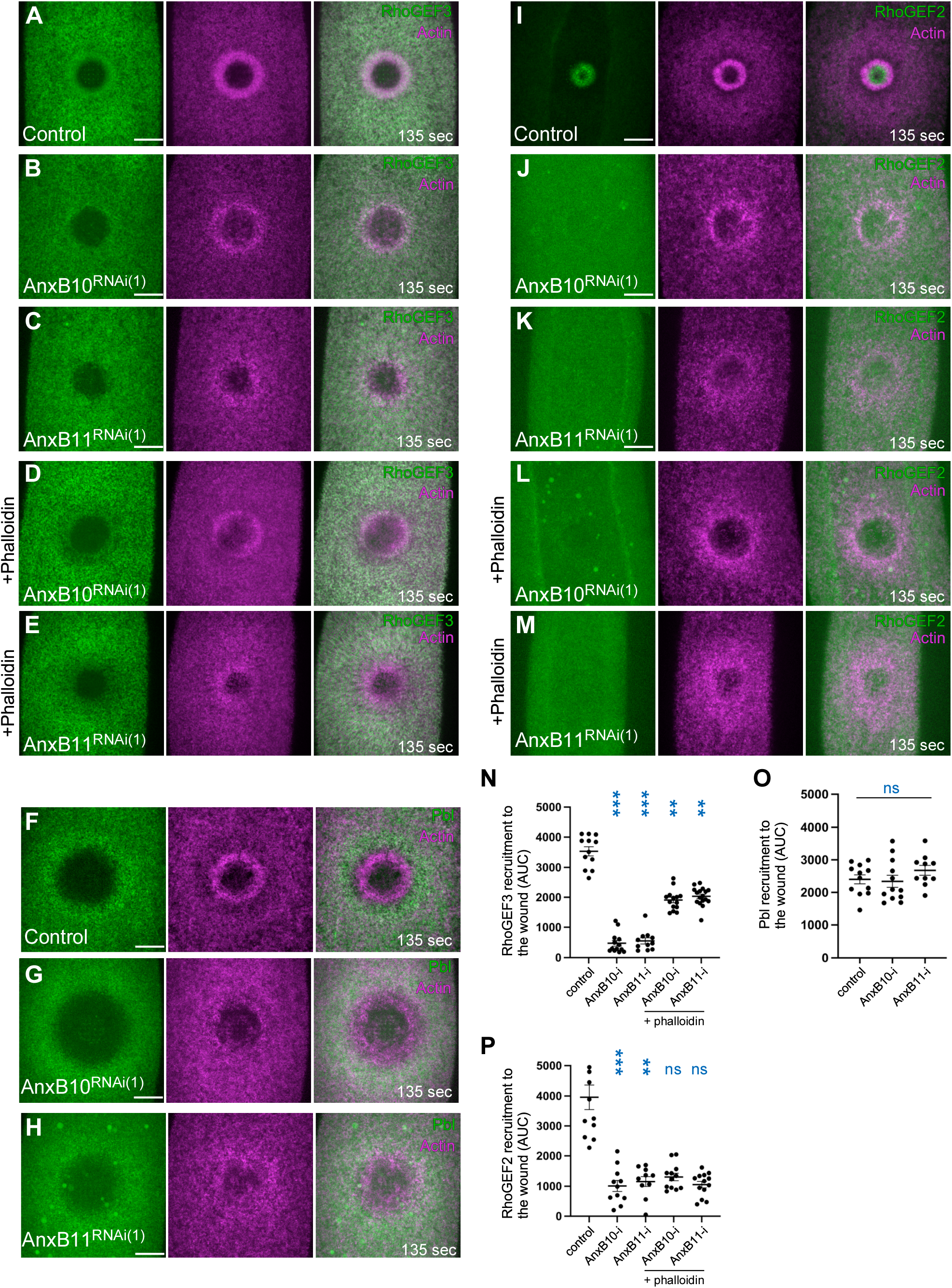
AnxB10 and AnxB11 are required for RhoGEF2 and RhoGEF3 recruitment to the wound. (A-E) Localization of sfGFP-RhoGEF3 along with actin reporter (sStMCA) in control (A), AnxB10^RNAi(1)^ (B), AnxB11^RNAi(1)^ (C), AnxB10^RNAi(1)^ + phalloidin (D), and AnxB11^RNAi(1)^ + phalloidin (E) backgrounds. (F-H) Localization of Pbl-eGFP along with actin reporter (sStMCA) in control (F), AnxB10^RNAi(1)^ (G), and AnxB11^RNAi(1)^ (H) backgrounds. (I-M) Localization of sfGFP-RhoGEF3 along with actin reporter (sStMCA) in control (I), AnxB10^RNAi(1)^ (J), AnxB11^RNAi(1)^ (K), AnxB10^RNAi(1)^ + phalloidin (L), and AnxB11^RNAi(1)^ + phalloidin (M) backgrounds. (N-P) Quantification of the area under the curve in each fluorescence intensity profile from individual embryos for RhoGEF3 (N), Pbl (O), and RhoGEF2 (P). ns, not significant. ANOVA tests were performed in N-P. n and time after wounding are indicated. Scale bar: 20 µm. Error bars represent ±SEM.

### Anx recruitment patterns to wounds are regulated by calcium dynamics rather than phospholipid membrane patterning

We next investigated how the rapid spatial recruitment patterns of Anxs to wounds are organized. Since Anxs are calcium-responding and phospholipid binding proteins (Blackwood and Ernst, 1990; Gerke et al., 2005; Lizarbe et al., 2013), we examined the possibilities that their specific recruitment patterns were in response to specific phospholipid patterns around the wound or in direct response to calcium dynamics at the wound. Previous studies showed that phospholipids such as PI(4, 5)P2, PI(3,4,5)P3, phosphatidylserine (PS), and phosphatidic acid (PA) in the plasma membrane are reorganized upon wounding in the tissue culture cell, *Xenopus* oocyte, and *Drosophila* cell wound models (Ashraf and Gerke, 2021; Nakamura et al., 2020; Vaughan et al., 2014). We first examined the specificity of the three *Drosophila* Anxs for binding phospholipids using membrane lipid strips and purified FLAG-tagged Anx proteins. In the absence of calcium, none of the Anxs bind to any of the phospholipids (Fig. S1C-D). In the presence of calcium, all Anxs bind to PE and PS, albeit with different preferences: AnxB9 binds equally to PE and PS, AnxB10 binds preferentially to PE, and AnxB11 binds preferentially to PS (Fig. S1C). In addition to these *in vitro* assays, available phospholipid biosensors, including one for PS, do not mimic Anx recruitment patterns around the wound (Fig. S1E-H; see Methods). Thus, Anxs recruitment patterns to wounds are not created by simply responding to phospholipid patterns around wounds. We next examined if calcium dynamics regulate Anx recruitment to wounds. Extracellular calcium influx is thought to be an initial wound repair start signal in many cell wound repair models (Andrews and Corrotte, 2018; Cooper and McNeil, 2015; Ebstrup et al., 2021; Moe et al., 2015; Nakamura et al., 2018). We previously found that while extracellular calcium is the primary calcium source in the *Drosophila* cell wound repair model, intracellular calcium sources are also required for robust wound repair (Nakamura et al., 2020). *Drosophila* embryos are surrounded by an impermeable vitelline membrane (Fig. 4A). Injection of the calcium chelator BAPTA (100 mM) into the perivitelline space (extracellular space between the embryo and vitelline membrane) removes almost all GCaMP signal, whereas injection of BAPTA into the embryo reduces the GCaMP signals around wounds (Fig. 4B, D-H). EGTA also chelates calcium, but it often exhibits different calcium responses compared to BAPTA due to their different working distances (Adler et al., 1991; Fakler and Adelman, 2008; Naraghi and Neher, 1997). Similar to BAPTA, injection of EGTA into the perivitelline space removes almost all GCaMP signal around wounds (Fig. 4C-E). Since BAPTA is more specific and has a higher affinity for Ca^2+^, we used BAPTA to alter calcium dynamics for the remaining experiments.

**Fig. 4.**
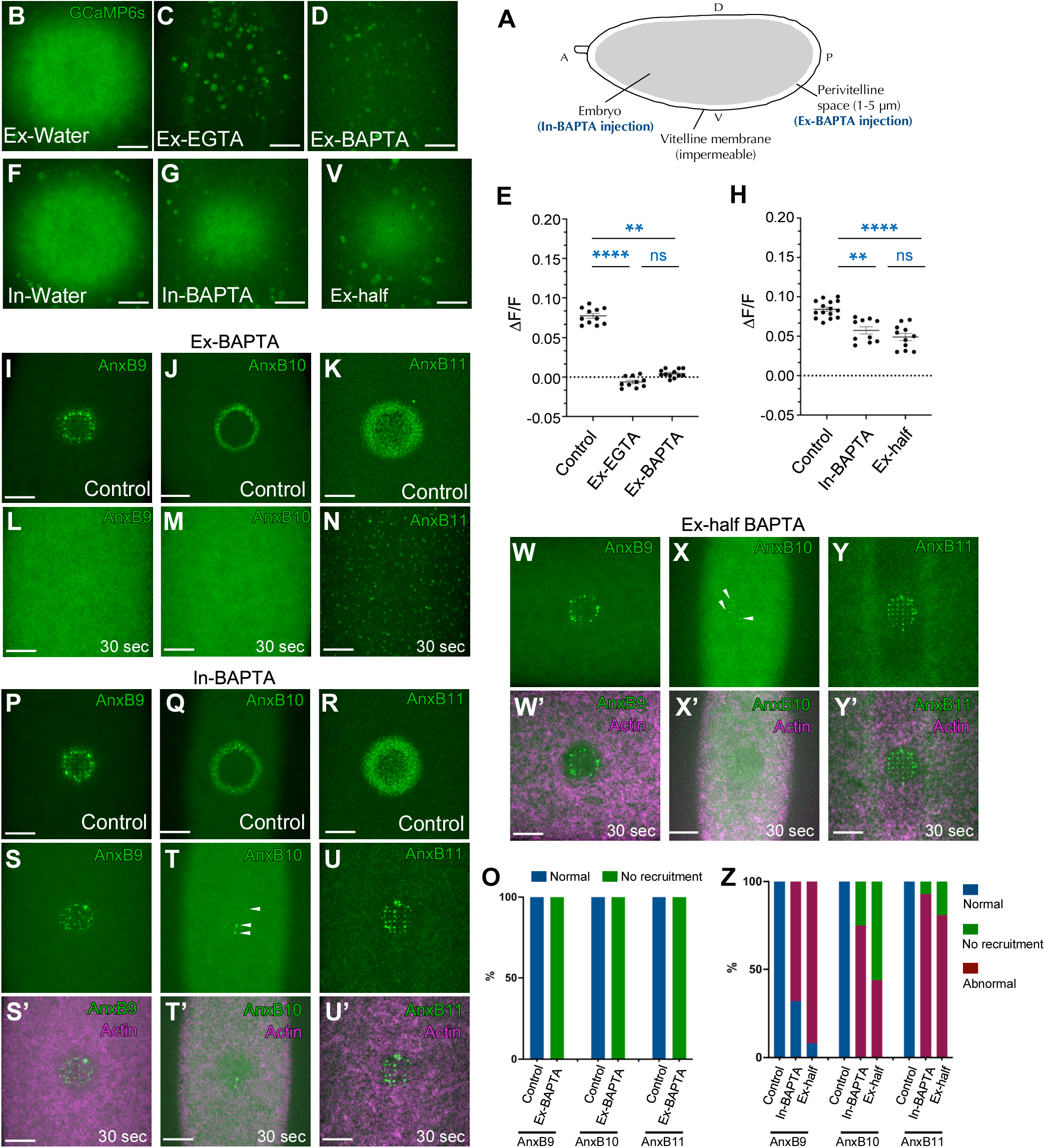
Calcium dynamics regulate Anx recruitment patterns to cell wounds. (A) Schemaric diagram showing the *Drosophila* embryo encapsulated by impermeable vitelline membrane. (B-D) GCaMP signals in NC4–6 staged *Drosophila* embryos with Ex-control water injection (B), Ex-EGTA (100 mM) injection (C), or Ex-BAPTA (100 mM) injection (D). (E) Quantification of GCaMP signals (ΔF/F) in the conditions of A-C. (F-G) GCaMP signals in NC4–6 staged *Drosophila* embryos with In-control water injection (F) or In-BAPTA (100 mM) injection (G). (H) Quantification of GCaMP signals (ΔF/F) in the conditions of E-F and V. (I-N) Localization of AnxB9, AnxB10, and AnxB11 in Ex-control (I-K) or Ex-BAPTA injection (J-N). (O) Quantification of Anx recruitment in control and Ex-BAPTA injection. (P-U’) Localization of AnxB9, AnxB10, and AnxB11 in In-control (P-R) and In-BAPTA injection (S-U’). (V) GCaMP signals in Ex-half BAPTA (50 mM) injection. (W-Y’) Localization of AnxB9, AnxB10, and AnxB11 in Ex-half BAPTA injection. (Z) Quantification of Anx recruitment in control, In-BAPTA injection, and Ex-half BAPTA injection. Time after wounding is indicated. Scale bar: 20 µm.

To test how the different calcium sources affect Anx recruitment patterns to wounds, we examined the recruitment of each Anx to wounds in the absence of extracellular (Ex-BAPTA) or intracellular (In-BAPTA) calcium. In the absence of extracellular calcium, none of the Anxs is recruited to wounds (Fig. 4I-N, O), which is consistent with previous studies using different models (Bittel et al., 2020; Demonbreun et al., 2016; Koerdt and Gerke, 2017). Surprisingly, when calcium is depleted intracellularly, all Anx recruitment patterns around wounds are different from that of the control (Fig. 4P-U’, Z). AnxB9 exhibits an asymmetric and punctate ring-like pattern (Fig. 4P, S-S’, Z). AnxB10 recruitment pattern is highly disrupted and exhibits a smaller number of puncta at the wound periphery (Fig. 4Q, T-T’, Z). AnxB11 is recruited inside the wound in a punctate pattern (Fig. 4R, U-U’, Z). To examine if the depletion of intracellular calcium or the total amount of calcium around wounds affects AnxB10 or AnxB11 recruitment patterns to wounds, we reduced extracellular calcium influx by injecting less BAPTA (50 mM; Ex-half) into the perivitelline space. We observed similar GCaMP signals around wounds as those detected upon the depletion of intracellular calcium (Fig. 4H, V). Intriguingly, Anx recruitment patterns to wounds in the reduced extracellular calcium influx are very similar to those observed upon depleting intracellular calcium (Fig. 4W-Y’, Z). Thus, rapid Anx recruitment patterns upon wounding are influenced by the quantity of the calcium influx. A previous study showed that temporal recruitment of Anx A1 and A6 to bacterial pore-forming wounds depends on their calcium sensitivity (Potez et al., 2011). The three *Drosophila* Anxs may similarly have different calcium sensitivities. Alternatively, while our current microscopy detects calcium signals as a uniform pattern around the wound within three seconds, rapid calcium diffusion may create gradient patterns. Development of more sensitive calcium reporters is needed to address this possibility.

### Depletion and reduced calcium influx from extracellular and intracellular sources affect cell wound repair differently

Since our results suggest that the quantity of calcium influx affects cell wound repair, we examined repair dynamics in the absence of extracellular or intracellular calcium (Fig. 5; Video 2). In the absence of extracellular calcium (Ex-BAPTA), no actin ring forms around the wound site and the wound does not close, albeit oscillations in wound size due to embryonic development were observed (Fig. 5A-C; Video 2). Thus, extracellular calcium is indispensable for repair in the *Drosophila* cell wound model. We expected that the wound in the absence of extracellular calcium would exhibit overexpansion since wound expansion and actin ring formation are correlated (Abreu-Blanco et al., 2011), and Anx knockdowns exhibit overexpansion phenotypes (Fig. 2L). Interestingly, wound expansion was not significantly different between control and extracellular BAPTA injected embryos (Fig 5D). Thus, wound expansion under the normal condition is likely due to the consequence of both positive and negative regulations.

**Fig. 5.**
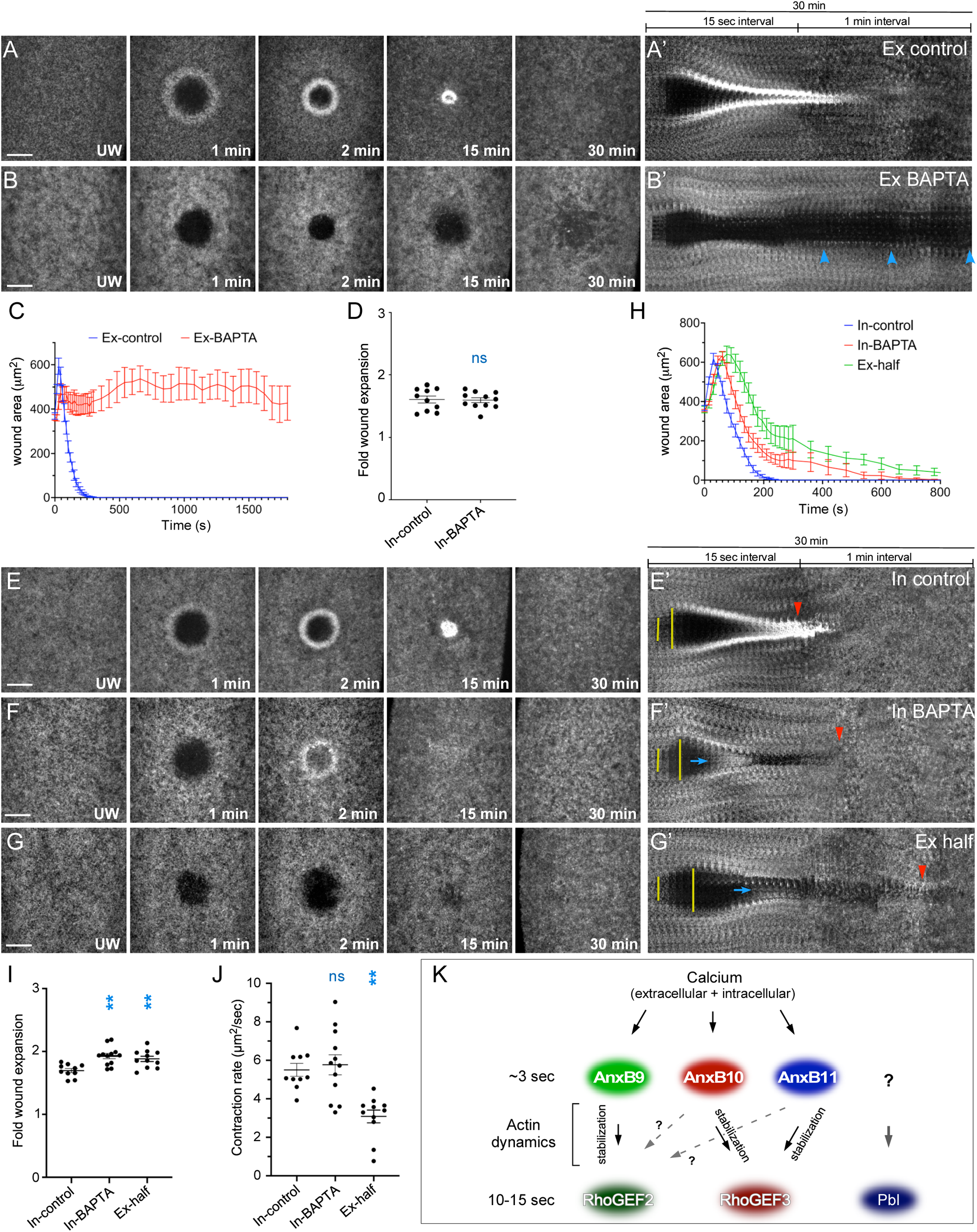
Lower calcium influx impairs cell wound repair. (A-B’) Actin dynamics (sGMCA or sStMCA) during cell wound repair in NC4–6 staged *Drosophila* embryos: Ex-control (A) and Ex-BAPTA (B). Failure of the wound to close is indicated by blue arrowheads. (C-D) Quantification of the wound area over time (C) and fold wound expansion (D) for Ex-control (n=10) and Ex-BAPTA (n=11). (E-G’) Actin dynamics (sGMCA or sStMCA) during cell wound repair in NC4–6 staged *Drosophila* embryos: In-control (E-E’), In-BAPTA (F-F’), and Ex-half (G-G’). Wound expansion is highlighted by yellow lines. Actin accumulation internal to the wound is indicated by blue arrows. Completion of wound closure is indicated by red arrowheads. (H-J) Quantification of the wound area over time (H), fold wound expansion (I), and contraction rate (J) for In-control (n=10), In-BAPTA (n=12), and Ex-half (n=11). (K) Schematic diagram summarizing the calcium > Anxs > actin > RhoGEFs pathways in response to cell wounds. Error bars represent ± SEM. Scale bar: 20 µm. Time after wounding is indicated. ANOVA tests were performed in D and I-J: ** *p*<0.01, ns is not significant.

In the absence of intracellular calcium (In-BAPTA), wounds exhibited overexpansion, delayed initiation of wound closure, actin accumulation inside the wound, a disrupted actin ring, and longer wound closure (Fig. 5E-F’, H-J; Video 2). Intriguingly, while Ex-half and In-BAPTA showed similar GCaMP signals, Ex-half had slower contraction rates and delayed wound onset/closure compared to In-BAPTA (Fig. 4H; Fig. 5E-E’, G-J; Video 2). These results suggest that, in addition to the amount of calcium signaling, the source of calcium has different effects on cell wound repair. It is possible that intracellular calcium release is slower than external influx during the first few seconds following wounding leading to more phenotypes occurring in Ex-half compared to those with In-BAPTA.

Previous studies and this study showed that cell wounds do not close in the absence of extracellular calcium (Bement et al., 1999; Idone et al., 2008; Jimenez et al., 2014; McNeil and Kirchhausen, 2005; McNeil and Terasaki, 2001; Talukder et al., 2020). In addition, calcium influx correlates with wound hole closure (Klenow et al., 2021) and the amount of calcium in the media affects survival rates after sonoporation, as well as actin accumulation levels at the wound (Talukder et al., 2020; Zhou et al., 2008), suggesting that calcium influx is also required for other repair events. We find that the quantity of calcium influx regulates the rapid recruitment patterns of Anxs around the wound, which in turn are required for the distinct recruitments of RhoGEF2 and RhoGEF3 mediated by actin dynamics including actin stabilization (Fig. 5K).

Since calcium works as a second messenger, dysregulation of calcium homeostasis is associated with different diseases, including different types of cancer, muscular dystrophy, and diabetes, which are also involved in cell wound repair (Ahn et al., 2017; Arruda and Hotamisligil, 2015; Cui et al., 2021; Levy, 1999; Monteith et al., 2017; Xu et al., 2018). Consistent with this, intracellular calcium levels from different sources in the disease conditions are known to enhance or impair cell wound repair in a context-dependent manner. For example, cancer cells have efficient cell wound repair in response to physical stresses during invasion processes (Bouvet et al., 2020; Ritter et al., 2022; Wirtz et al., 2011), whereas diseased cells from muscular dystrophy or diabetes backgrounds exhibit poor cell wound repair (Bansal et al., 2003; Cai et al., 2009; Chandra et al., 2019; Cheng et al., 2014; Mareedu et al., 2021; Quattrocelli et al., 2017; Tsurumi et al., 2019).

We previously showed that insulin signaling is activated upon cell wounding to regulate actin dynamics (Nakamura et al., 2020). Intriguingly, our results showing that Ex-half and In-BAPTA impair cell wound repair differently suggest that cells can sense different sources of calcium to activate distinct downstream pathways. Different sources of calcium might affect dysregulation of calcium homeostasis differentially, leading to different cell wound repair efficiency in disease conditions. Future challenges include investigating how calcium influx is able to rapidly regulate protein recruitment to the wound periphery and how sources of calcium affect cell wound repair differently, thereby impacting how a cell responds to an injury and initiates repair processes in different conditions.

## ACKNOWLEDGEMENTS

We thank Du Guangwei, Tony Cooke, the Bloomington Stock Center, the Kyoto Stock Center, the Harvard Transgenic RNAi Project, FlyBase, the Fred Hutch/Leica Center of Excellence, and the Drosophila Genome Resource Center for advice, microscopes, antibodies, DNAs, flies, and other reagents used in this study.

## Author Contributions

MN and SMP designed the experiments and analyzed the resulting data. MN performed the experiments. MN and SMP wrote the manuscript.

## Abbreviations List

Anx: Annexin
BSA: Bovine serum albumin
GFP: Green fluorescent protein
NC: Nuclear cycle
PA: Phosphatidic acid
Pbl: Pebble
PS: Phosphatidylserine
RhoGAP: Rho GTPase activating protein
RhoGEF: Rho guanine nucleotide exchange factor
sfGFP: super folder green fluorescent protein
sGMCA: spaghetti squash driven, GFP, moesin-α-helical-coiled and actin binding site
sqh: spaghetti squash
sStMCA: spaghetti squash driven, mScarlet, moesin-α-helical-coiled and actin binding site
StFP: mScarlet fluorescent protein

## MATERIALS AND METHODS

**Reagents used in this study are described in Table S1.**

### Fly stocks and genetics

Flies were cultured and crossed at 25°C on yeast-cornmeal-molasses-malt medium. Flies used in this study are described in Table S1. All fly stocks were treated with tetracycline, then tested by PCR to ensure that they did not harbor Wolbachia.

To knockdown genes, RNAi lines were driven maternally using the GAL4-UAS system (Brand and Perrimon, 1993) with P{matalpha4-GAL-VP16}V37 and used two independent RNAi lines each for AnxB10 and AnxB11 (Table S1). Localization patterns and mutant analyses were performed at least twice from independent genetic crosses and ≥10 embryos were examined unless otherwise noted. Images representing the average phenotype were selected for figures.

### Generation of fluorescently tagged Annexins and lipid biosensors

RNAi lines for AnxB11 were generated using the method previously described (Ni et al., 2011). Two oligos were annealed and cloned into pWALIUM22 (DGRC_1473). pWALIUM22-AnxB11 RNAi(1) and RNAi(2) were injected into M{3xP3-RFP.attP}ZH-86Fb (BDSC_24749).

To generate the StFP knock-in AnxB10, we used CRIMIC technology (Lee et al., 2018). We replaced the GAL4 cassette in AnxB10[CR00582-TG4.0] allele with StFP.

To generate sqh-sfGFP-AnxB10, the AnxB10 ORF was amplified from BDGP clone LD25605 and fused 5’ to sfGFP. The resulting sfGFP-AnxB10 fusion was cloned into pSqh5’+3’UTR (MT) as a 5’ StuI-3’ XbaI fragment.

To generate AnxB11-StFP, a genomic region of AnxB11, including 5kb upstream from the start codon and 2.2 kb downstream from the stop codon, was amplified and cloned into pCasper4 with StFP inserted just before the AnxB11 start codon.

To generate Ubi-StFP-8PM, StFP-8PM was amplified by adding prenylated octavalent peptides (+8pre: ARDGRRRRRRARARCVIM; (Eisenberg et al., 2021) to C-terminus of StFP. StFP-8PM was cloned into pUbi5’+3’UTR as a 5’ StuI-3’ XbaI fragment. StFP-8PM binds to plasma membrane electrostatic properties dependently and gives a cleaner label than general membrane reporters/dyes such as FM-4-64.

To generate the PS biosensor, Lact-C2 was amplified from Lact-C2-GFP (Addgene #22852) and fused 5’ to StFP. The resulting Lact-C2-StFP fusion was cloned into UASz as a 5’ KpnI and 3’ XbaI fragment.

To generate the PA biosensor, the PASS sequence was amplified from a GFP-PASS plasmid (provided by G. Du, University of Texas Health Science Center at Houston, Houston, Texas; (Zhang et al., 2014)) and fused 5’ to sfGFP. The resulting sfGFP-PASS fusion was cloned into UASz as a 5’ KpnI and 3’ XbaI fragment.

To generate the PI(3,5)P2 biosensor, ML1Nwt-eGFP was amplified from ML1Nwt_in_pEGFP-C1 (Addgene #67797; (Li et al., 2013)) and cloned into pSqh5’+3’UTR as a 5′ StuI-3′ XbaI fragment.

To generate transgenic flies, each construct (500 µg/ml) was injected along with the pTURBO helper plasmid (100 µg/ml) into isogenic w1118 flies as previously described (Spradling, 1986).

### Embryo handling and preparation

Nuclear cycle (NC) 4-6 *Drosophila* embryos were collected from 0-30 min at room temperature (22°C). Embryos were hand dechorionated, placed onto No. 1.5 coverslips coated with glue, and covered with Series 700 halocarbon oil (Halocarbon Products Corp).

### Laser wounding

All wounds were generated with a pulsed nitrogen N2 Micropoint laser (Andor Technology Ltd., Concord, MA, USA) tuned to 435 nm and focused on the cortical surface of the embryo. A region of interest was selected in the lateral midsection of the embryo and ablation was controlled by MetaMorph. On average, ablation time was less than 3s, and time-lapse imaging was initiated immediately. Occasionally, a faint grid pattern of fluorescent dots is visible at the center of wounds that arises from damage to the vitelline membrane that covers embryos.

### Drug injections

Pharmacological inhibitors were injected into NC4–6 staged *Drosophila* embryos, incubated at room temperature (22°C) for 5 min, and then subjected to laser wounding. The following inhibitors were used: phalloidin (100 µg/ml; Thermo Fisher Scientific), BAPTA (100 mM or 50 mM; Invitrogen), and EGTA (100 mM; Sigma-Aldrich). Phalloidin was prepared in injection buffer (5 mM KCl, 0.1 mM NaP, pH 6.8). BAPTA and EGTA were prepared in water.

### Microscopy

All imaging was performed at room temperature (22°C). The following microscope was used: Revolution WD systems (Andor Technology Ltd., Concord, MA, USA) mounted on a Leica DMi8 (Leica Microsystems Inc., Buffalo Grove, IL, USA) with a 63x/1.4 NA objective lens and controlled by MetaMorph software. Images and videos were acquired with 488 nm and 561 nm, using an Andor iXon Ultra 897 or 888 EMCCD cameras (Andor Technology Ltd., Concord, MA, USA). All images for cell wound repair were 12-20 µm stacks/0.25 µm steps. To examine each Anx recruitment to the wound without other reporters, images were acquired every 5 sec. To examine Anx recruitment relative to actin and other Anxs, images were acquired every 15 sec with dual camera mode using two Andor iXon 888. To examine calcium dynamics using GCaMP6s, images were acquired every 15 sec. To examine RNAi knockdown phenotypes, images were acquired every 30 sec for 15 min and then every 60 sec for 25 min. To examine RhoGEF recruitment to the wound, images were acquired every 45 sec.

### Image processing, analysis, and quantification

All images were analyzed with Fiji (Schindelin et al., 2012) and Matlab (Mathworks). Measurements of wound area were done manually. To generate xy kymographs, all time-lapse xy images were cropped to 5.5 µm x 98.1 µm and then each cropped image was lined up. For fluorescent line plots, the mean fluorescence profile intensities were calculated from 51 equally spaced radial profiles anchored at the center of the wound, swept from 0° to 180° (Hui et al). Radial profiles of diameter 301 pixels were used. To normalize the signal intensity in ≥10 embryos, the intensity in each pixel was divided by max intensity in each embryo. The lines represent the averaged fluorescent intensity from ≥10 embryos and gray area is the 95% confidence interval.

Quantification of the width and average intensity of actin ring, wound expansion, and closure rate was performed as follows: the width of actin ring was calculated with two measurements, the ferret diameters of the outer and inner edge of actin ring at 120 sec post-wounding. Using these measurements, the width of actin ring was calculated with (outer feret diameter – inner feret dimeter)/2. The average intensity of actin ring was calculated with two measurements. Instead of measuring ferret diameters, we measured area and integrated intensity in same regions as described in ring width. Using these measurements, the average intensity in the actin ring was calculated with (outer integrated intensity - inner integrated intensity)/(outer area - inner area). To calculate relative intensity for unwounded (UW) time point, average intensity at UW was measured with 50×50 pixels at the center of embryos and then averaged intensity of actin ring at each timepoint was divided by average intensity of UW. Wound expansion was calculated with max wound area/initial wound size. Closure rate was calculated with two time points, one is t_max_ that is the time of reaching maximum wound area, the other is t<half that is the time of reaching 50-35% size of max wound since the slope of wound area curve changes after t<half. Using these time points, average speed was calculated with (wound area at t_max_ – wound area at t<half)/t_max_-t<half. To quantify RhoGEF recruitment to wounds, we subtracted the fluorescent intensity of the pre-wounding time point from the 135 sec post-wounding image. We then measured the averaged fluorescent intensity from radial line plots (described above) in the subtracted image. Line profiles were plotted and area under the curve (AUC) was measured (Nakamura et al., 2017). Generation of all graphs and a one-way ANOVA test were performed with Prism 8 (GraphPad Software Inc.) or Matlab (Mathworks).

### Protein expression

AnxB9, AnxB10 and AnxB11 cDNAs were amplified as 5’ BamHI-3’ NotI fragments from cDNA clones (DGRC) and then cloned into a double tag pGEX-dt vector (GST and His; (Liu et al., 2009) with FLAG tag at the N-terminus of Anxs. Protein expression was performed as previously described (Rosales-Nieves et al., 2006). Cells were lysed by sonication in lysis buffer (50 mM Tris pH 7.6, 100 mM NaCl, 1 mM EGTA, 5% Glycerol, 1mM DTT) with 1% Triton-X, 50 mM imidazole, and Complete protease inhibitor tablets (ThermoFisher). Lysates were centrifuged at 10,000 g for 30 min and the supernatants were coupled to Fastflow nickel-sepharose (GE) for 3 hours at 4°C. The matrix was washed three times with lysis buffer with 50 mM imidazole and eluted by lysis buffer with 1 M imidazole. All His elutions were coupled to glutathione-sepharose 4B (GE) for 3 hours at 4°C, washed with lysis buffer, and then eluted with HRV 3C protease (ThermoFisher). All proteins were dialyzed into storage buffer (50 mM Tris pH 7.6, 100 mM NaCl, 5% Glycerol, 1mM DTT) and then flash frozen.

### Lipid protein interaction assay

PIP strip membranes (Invitrogen) were blocked in blocking buffer (TBS: 20 mM Tris pH 8.0 + 150 mM NaCl, with 0.1% Tween-20 and 1% non-fat milk) for one hour at RT and incubated with purified Anx proteins (0.5 µg/ml) and 100 µM CaCl_2_ or 100 µM EGTA for one hour at RT. The membranes were washed three times with TBS-T (TBS with 0.1% Tween-20) and bound Anxs were detected by immunoblotting using Rat anti-FLAG antibody (1:5000; Biolegend) and ECL (ThermoFisher).

### qPCR

Total RNA was obtained from 100 embryos (0-30 min old) using TRIzol (Invitrogen). 1 µg of total RNA was used for reverse transcription with the iScript™ gDNA Clear cDNA Synthesis Kit (Bio-Rad). RT-PCR analysis was performed using the iTaq™ Universal SYBR® Green Supermix (Bio-Rad) with two individual parent sets and two technical replicates on the CFX96TM Real Time PCR Detection System (Bio-Rad). RpL32 was used as a reference gene. The % knockdown was calculated using the ΔΔCq calculation method compared with control (vermilion knockdown). Same primer sets RpL32 was used in previously described (Nakamura et al., 2020).

### Statistical analysis

All statistical analysis was done using Prism 8 (GraphPad, San Diego, CA). Gene knockdowns were compared to the appropriate control, and statistical significance was calculated using a one-way ANOVA test with *P*<0.01 considered significant.

## SUPPLEMENTARY FIGURE LEGENDS

**Fig. S1.**
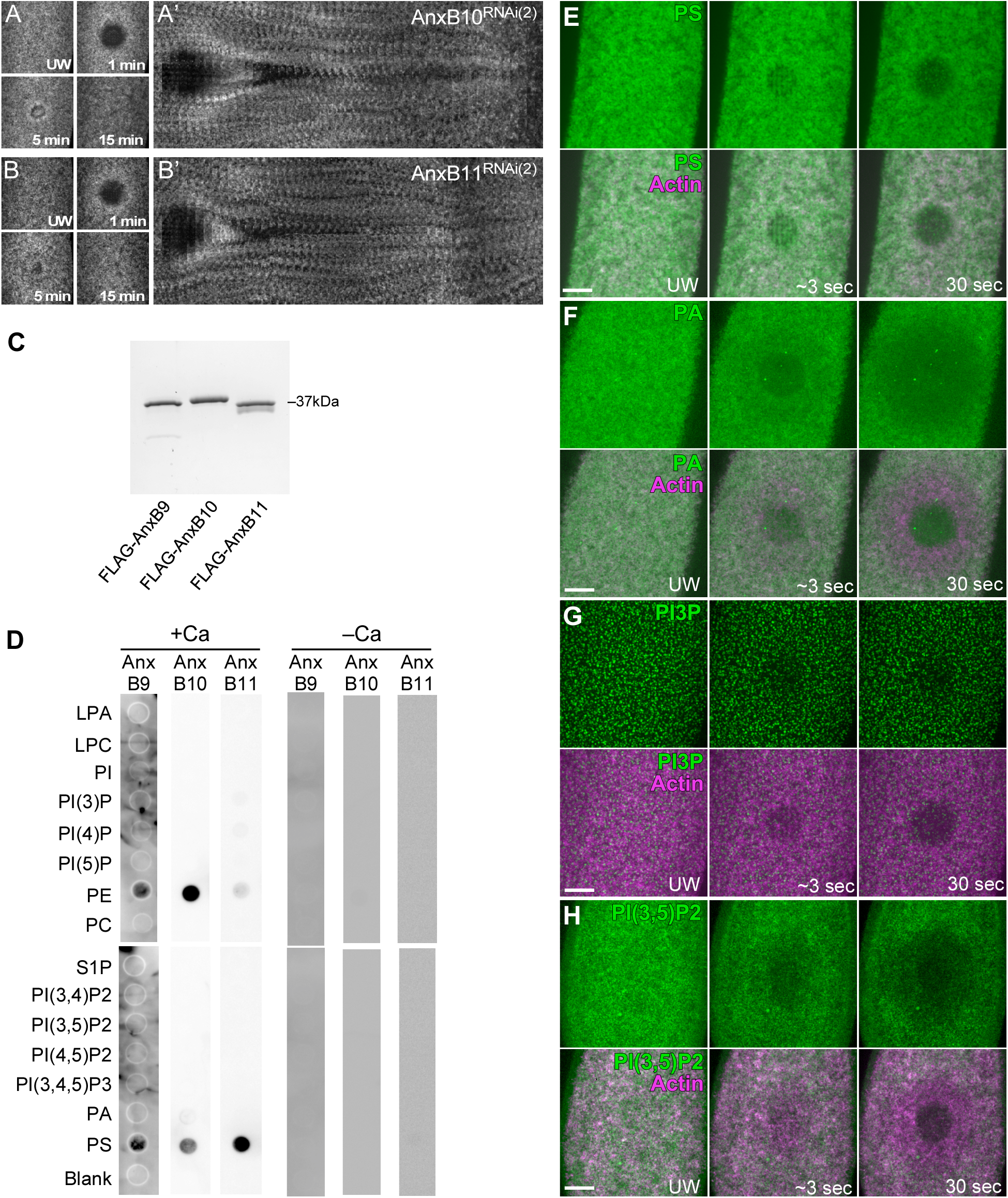
AnxB10 and AnxB11 RNAi phenotypes and membrane lipids in cell wound repair. (A-B’) Actin dynamics (sGMCA) during cell wound repair in NC4–6 staged *Drosophila* embryos: AnxB10^RNAi(2)^ (A-A’) and AnxB11^RNAi(2)^ (B-B’). (C) Coomassie stained gel showing the purified Anx proteins used in this study. (D) PIP strip assays. Purified recombinant FLAG-AnxB9, FLAG-AnxB10, and FLAG-AnxB11 proteins were incubated with a PIP strip +/− Ca^2+^. (E-H) Localization of PS (E), PA (F), PI3P (G), and PI(3,5)P2 (H) lipid reporters along with an actin reporter (sGMCA or sStMCA). Time after wounding is indicated. Scale bar: 20 µm.Time after wounding is indicated. Scale bar: 20 µm.

## SUPPLEMENTARY VIDEO LEGENDS

**Video 1.** Three *Drosophila* Anxs have non-redundant roles during cell wound repair (A-G) Time-lapse confocal xy images and fluorescence intensity (arbitrary units) profiles across the wound area from Drosophila NC4-6 staged embryos expressing an actin reporter (sGMCA, magenta) in: control (A), AnxB9-i (B), AnxB10-i(1) (C), AnxB10-i(2) (D), AnxB11i-(1) (E), AnxB11-i(2) (F), and triple Anx-i (G). (H) Time-lapse confocal xy images and fluorescence intensity (arbitrary units) profiles across the wound area from Drosophila NC4-6 staged embryos co-expressing an actin reporter (sGMCA, green) along with StFP-8PM (magenta) in triple Anx knockdowns. Time post-wounding is indicated. UW: unwounded.

**Video 2.** Depletion of extra- and intra-cellular calcium impairs cell wound repair differently (A-E) Time-lapse confocal xy images and fluorescence intensity (arbitrary units) profiles across the wound area from Drosophila NC4-6 staged embryos expressing an actin reporter (sGMCA) in: Ex-control (A), Ex-BAPTA (B), In-control (C), In-BAPTA (D), and Ex-half (E). Time post-wounding is indicated. UW: unwounded.

**Table S1.**
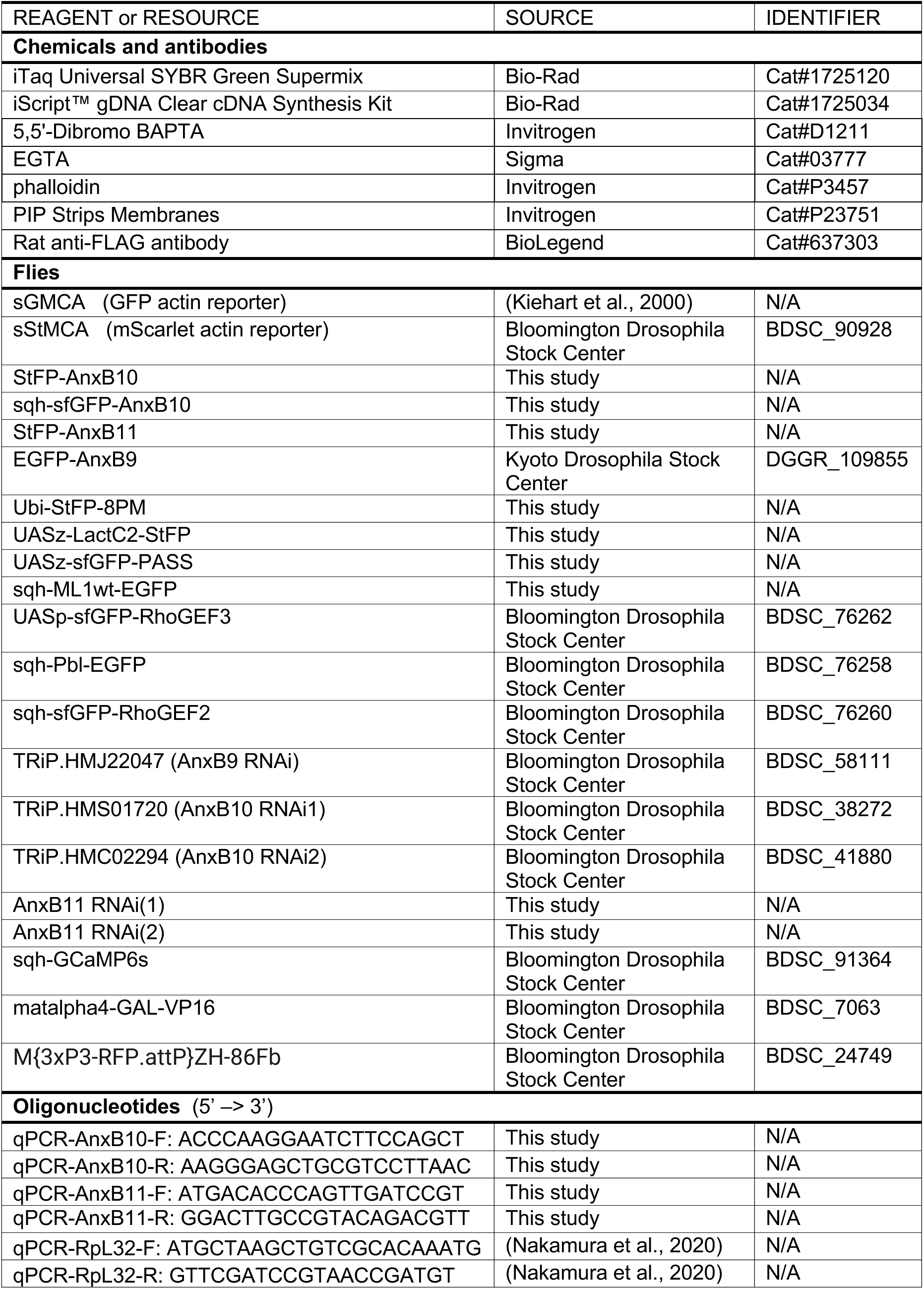
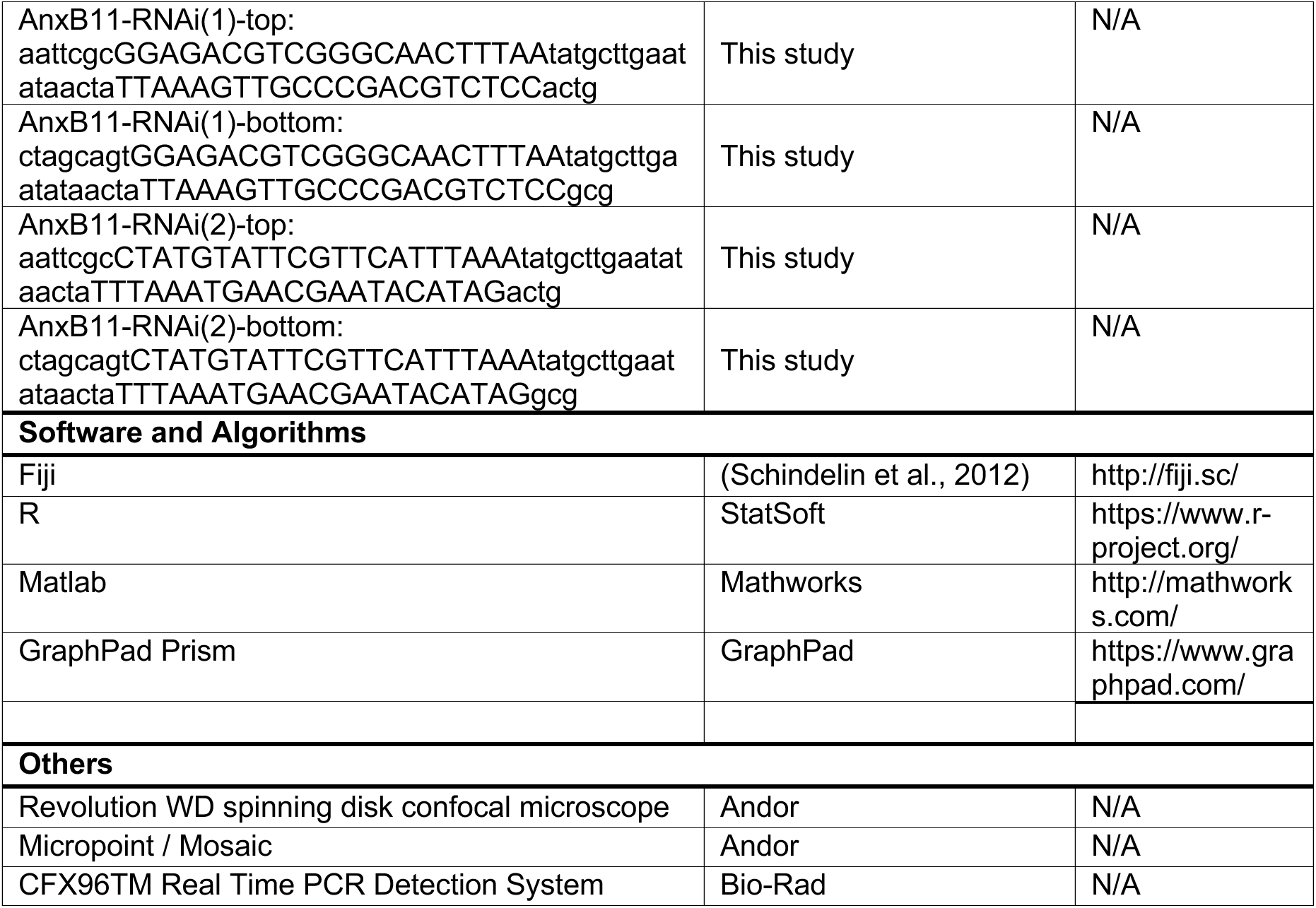
Flies and reagents used in this study.

## REFERENCES

Abreu-Blanco, M.T., J.M. Verboon, and S.M. Parkhurst. 2011. Cell wound repair in Drosophila occurs through three distinct phases of membrane and cytoskeletal remodeling. J Cell Biol. 193:455–464.

Adler, E.M., G.J. Augustine, S.N. Duffy, and M.P. Charlton. 1991. Alien intracellular calcium chelators attenuate neurotransmitter release at the squid giant synapse. J Neurosci. 11:1496–1507.

Ahn, C., J.H. Kang, and E.B. Jeung. 2017. Calcium homeostasis in diabetes mellitus. J Vet Sci. 18:261–266.

Andrews, N.W., and M. Corrotte. 2018. Plasma membrane repair. Curr Biol. 28:R392–R397.

Arruda, A.P., and G.S. Hotamisligil. 2015. Calcium Homeostasis and Organelle Function in the Pathogenesis of Obesity and Diabetes. Cell Metab. 22:381–397.

Ashraf, A.P.K., and V. Gerke. 2021. Plasma membrane wound repair is characterized by extensive membrane lipid and protein rearrangements in vascular endothelial cells. Biochim Biophys Acta Mol Cell Res. 1868:118991.

Babbin, B.A., C.A. Parkos, K.J. Mandell, L.M. Winfree, O. Laur, A.I. Ivanov, and A. Nusrat. 2007. Annexin 2 regulates intestinal epithelial cell spreading and wound closure through Rho-related signaling. The American journal of pathology. 170:951–966.

Bansal, D., K. Miyake, S.S. Vogel, S. Groh, C.C. Chen, R. Williamson, P.L. McNeil, and K.P. Campbell. 2003. Defective membrane repair in dysferlin-deficient muscular dystrophy. Nature. 423:168–172.

Bement, W.M., C.A. Mandato, and M.N. Kirsch. 1999. Wound-induced assembly and closure of an actomyosin purse string in Xenopus oocytes. Current biology : CB. 9:579–587.

Benaud, C., G. Le Dez, S. Mironov, F. Galli, D. Reboutier, and C. Prigent. 2015. Annexin A2 is required for the early steps of cytokinesis. EMBO Rep. 16:481–489.

Benaud, C., and C. Prigent. 2016. Annexin A2: A new player in mitosis. In Cell Cycle. Vol. 15. 9–10.

Bittel, D.C., G. Chandra, L.M.S. Tirunagri, A.B. Deora, S. Medikayala, L. Scheffer, A. Defour, and J.K. Jaiswal. 2020. Annexin A2 Mediates Dysferlin Accumulation and Muscle Cell Membrane Repair. Cells. 9:1919.

Blackwood, R.A., and J.D. Ernst. 1990. Characterization of Ca2(+)-dependent phospholipid binding, vesicle aggregation and membrane fusion by annexins. Biochem J. 266:195–200.

Bouter, A., C. Gounou, R. Berat, S. Tan, B. Gallois, T. Granier, B.L. d’Estaintot, E. Poschl, B. Brachvogel, and A.R. Brisson. 2011. Annexin-A5 assembled into two-dimensional arrays promotes cell membrane repair. Nat Commun. 2:270.

Bouvet, F., M. Ros, E. Bonedeau, C. Croissant, L. Frelin, F. Saltel, V. Moreau, and A. Bouter. 2020. Defective membrane repair machinery impairs survival of invasive cancer cells. Sci Rep. 10:21821.

Boye, T.L., K. Maeda, W. Pezeshkian, S.L. Sønder, S.C. Haeger, V. Gerke, A.C. Simonsen, and J. Nylandsted. 2017. Annexin A4 and A6 induce membrane curvature and constriction during cell membrane repair. Nature Communications. 8:1623.

Brand, A.H., and N. Perrimon. 1993. Targeted gene expression as a means of altering cell fates and generating dominant phenotypes. Development. 118:401–415.

Cai, C., N. Weisleder, J.K. Ko, S. Komazaki, Y. Sunada, M. Nishi, H. Takeshima, and J. Ma. 2009. Membrane repair defects in muscular dystrophy are linked to altered interaction between MG53, caveolin-3, and dysferlin. J Biol Chem. 284:15894–15902.

Carmeille, R., F. Bouvet, S. Tan, C. Croissant, C. Gounou, K. Mamchaoui, V. Mouly, A.R. Brisson, and A. Bouter. 2016. Membrane repair of human skeletal muscle cells requires Annexin-A5. Biochim Biophys Acta. 1863:2267–2279.

Chandra, G., A. Defour, K. Mamchoui, K. Pandey, S. Mishra, V. Mouly, S. Sreetama, M. Mahad Ahmad, I. Mahjneh, H. Morizono, N. Pattabiraman, A.K. Menon, and J.K. Jaiswal. 2019. Dysregulated calcium homeostasis prevents plasma membrane repair in Anoctamin 5/TMEM16E-deficient patient muscle cells. Cell Death Discov. 5:118.

Cheng, X., X. Zhang, Q. Gao, M. Ali Samie, M. Azar, W.L. Tsang, L. Dong, N. Sahoo, X. Li, Y. Zhuo, A.G. Garrity, X. Wang, M. Ferrer, J. Dowling, L. Xu, R. Han, and H. Xu. 2014. The intracellular Ca(2)(+) channel MCOLN1 is required for sarcolemma repair to prevent muscular dystrophy. Nat Med. 20:1187–1192.

Cheng, X., X. Zhang, L. Yu, and H. Xu. 2015. Calcium signaling in membrane repair. Semin Cell Dev Biol. 45:24–31.

Cooper, S.T., and P.L. McNeil. 2015. Membrane Repair: Mechanisms and Pathophysiology. Physiol Rev. 95:1205–1240.

Croissant, C., C. Gounou, F. Bouvet, S. Tan, and A. Bouter. 2020. Annexin-A6 in Membrane Repair of Human Skeletal Muscle Cell: A Role in the Cap Subdomain. Cells. 9:1742.

Cui, C., Y. Zhang, G. Liu, S. Zhang, J. Zhang, and X. Wang. 2021. Advances in the study of cancer metastasis and calcium signaling as potential therapeutic targets. Explor Target Antitumor Ther. 2:266–291.

Davenport, N.R., K.J. Sonnemann, K.W. Eliceiri, and W.M. Bement. 2016. Membrane dynamics during cellular wound repair. Mol Biol Cell. 27:2272–2285.

Demonbreun, A.R., E. Bogdanovic, L.A. Vaught, N.L. Reiser, K.S. Fallon, A.M. Long, C.C. Oosterbaan, M. Hadhazy, P.G.T. Page, P.R.B. Joseph, G. Cowen, A.M. Telenson, A. Khatri, K.R. Sadleir, R. Vassar, and E.M. McNally. 2022. A conserved annexin A6–mediated membrane repair mechanism in muscle, heart, and nerve.

Demonbreun, A.R., M. Quattrocelli, D.Y. Barefield, M.V. Allen, K.E. Swanson, and E.M. McNally. 2016. An actin-dependent annexin complex mediates plasma membrane repair in muscle. J Cell Biol. 213:705–718.

Ebstrup, M.L., C. Dias, A.S.B. Heitmann, S.L. Sønder, and J. Nylandsted. 2021. Actin Cytoskeletal Dynamics in Single-Cell Wound Repair. International Journal of Molecular Sciences. 22:10886.

Eisenberg, S., E. Haimov, G.F.W. Walpole, J. Plumb, M.M. Kozlov, and S. Grinstein. 2021. Mapping the electrostatic profiles of cellular membranes. Mol Biol Cell. 32:301–310.

Fakler, B., and J.P. Adelman. 2008. Control of K(Ca) channels by calcium nano/microdomains. Neuron. 59:873–881.

Gabel, M., F. Delavoie, V. Demais, C. Royer, Y. Bailly, N. Vitale, M.F. Bader, and S. Chasserot-Golaz. 2015. Annexin A2-dependent actin bundling promotes secretory granule docking to the plasma membrane and exocytosis. J Cell Biol. 210:785–800.

Gerke, V., C.E. Creutz, and S.E. Moss. 2005. Annexins: linking Ca2+ signalling to membrane dynamics. Nat Rev Mol Cell Biol. 6:449–461.

Glenney, J.R., Jr., B. Tack, and M.A. Powell. 1987. Calpactins: two distinct Ca++-regulated phospholipid- and actin-binding proteins isolated from lung and placenta. J Cell Biol. 104:503–511.

Hayes, M.J., U. Rescher, V. Gerke, and S.E. Moss. 2004. Annexin-actin interactions. Traffic. 5:571–576.

Hosoya, H., R. Kobayashi, S. Tsukita, and F. Matsumura. 1992. Ca(2+)-regulated actin and phospholipid binding protein (68 kD-protein) from bovine liver: identification as a homologue for annexin VI and intracellular localization. Cell Motil Cytoskeleton. 22:200–210.

Hui, J., V. Stjepic, M. Nakamura, and S.M. Parkhurst. 2022. Wrangling Actin Assemblies: Actin Ring Dynamics during Cell Wound Repair. Cells. 11.

Idone, V., C. Tam, J.W. Goss, D. Toomre, M. Pypaert, and N.W. Andrews. 2008. Repair of injured plasma membrane by rapid Ca2+-dependent endocytosis. J Cell Biol. 180:905–914.

Ikebuchi, N.W., and D.M. Waisman. 1990. Calcium-dependent regulation of actin filament bundling by lipocortin-85. J Biol Chem. 265:3392–3400.

Jimenez, A.J., P. Maiuri, J. Lafaurie-Janvore, S. Divoux, M. Piel, and F. Perez. 2014. ESCRT Machinery Is Required for Plasma Membrane Repair.pdf. Science (New York, N.Y.). 343:1247136–1247136.

Kiehart, D.P., C.G. Galbraith, K.A. Edwards, W.L. Rickoll, and R.A. Montague. 2000. Multiple Forces Contribute to Cell Sheet Morphogenesis for Dorsal Closure in Drosophila. The Journal of Cell Biology. 149:471–490.

Klenow, M.B., A.S.B. Heitmann, J. Nylandsted, and A.C. Simonsen. 2021. Timescale of hole closure during plasma membrane repair estimated by calcium imaging and numerical modeling. Sci Rep. 11:4226.

Koerdt, S.N., and V. Gerke. 2017. Annexin A2 is involved in Ca(2+)-dependent plasma membrane repair in primary human endothelial cells. Biochim Biophys Acta. 1864:1046–1053.

Lee, P.T., J. Zirin, O. Kanca, W.W. Lin, K.L. Schulze, D. Li-Kroeger, R. Tao, C. Devereaux, Y. Hu, V. Chung, Y. Fang, Y. He, H. Pan, M. Ge, Z. Zuo, B.E. Housden, S.E. Mohr, S. Yamamoto, R.W. Levis, A.C. Spradling, N. Perrimon, and H.J. Bellen. 2018. A gene-specific T2A-GAL4 library for Drosophila. Elife. 7.

Lennon, N.J., A. Kho, B.J. Bacskai, S.L. Perlmutter, B.T. Hyman, and R.H. Brown. 2003. Dysferlin interacts with annexins A1 and A2 and mediates sarcolemmal wound-healing. The Journal of biological chemistry. 278:50466–50473.

Levy, J. 1999. Abnormal cell calcium homeostasis in type 2 diabetes mellitus: a new look on old disease. Endocrine. 10:1–6.

Li, X., X. Wang, X. Zhang, M. Zhao, W.L. Tsang, Y. Zhang, R.G.W. Yau, L.S. Weisman, and H. Xu. 2013. Genetically encoded fluorescent probe to visualize intracellular phosphatidylinositol 3,5-bisphosphate localization and dynamics. In Proc Natl Acad Sci U S A. Vol. 110. 21165–21170.

Liu, R., M.T. Abreu-Blanco, K.C. Barry, E.V. Linardopoulou, G.E. Osborn, and S.M. Parkhurst. 2009. Wash functions downstream of Rho and links linear and branched actin nucleation factors. Development. 136:2849–2860.

Lizarbe, M., J. Barrasa, N. Olmo, F. Gavilanes, and J. Turnay. 2013. Annexin-Phospholipid Interactions. Functional Implications. International Journal of Molecular Sciences. 14:2652–2683.

Mareedu, S., E.D. Million, D. Duan, and G.J. Babu. 2021. Abnormal Calcium Handling in Duchenne Muscular Dystrophy: Mechanisms and Potential Therapies. Front Physiol. 12:647010.

McNeil, A.K., U. Rescher, V. Gerke, and P.L. McNeil. 2006. Requirement for annexin A1 in plasma membrane repair. The Journal of biological chemistry. 281:35202–35207.

McNeil, P.L., and T. Kirchhausen. 2005. An emergency response team for membrane repair. Nature reviews. Molecular cell biology. 6:499–505.

McNeil, P.L., and M. Terasaki. 2001. Coping with the inevitable: how cells repair a torn surface membrane. Nature cell biology. 3:E124–E129.

Moe, A., A.E. Golding, and W.M. Bement. 2015. Cell healing: Calcium, repair and regeneration. Seminars in Cell & Developmental Biology.

Monteith, G.R., N. Prevarskaya, and S.J. Roberts-Thomson. 2017. The calcium–cancer signalling nexus. Nature Reviews Cancer. 17:373–380.

Morel, E., R.G. Parton, and J. Gruenberg. 2009. Annexin A2-dependent polymerization of actin mediates endosome biogenesis. Developmental cell. 16:445–457.

Nakamura, M., A.N.M. Dominguez, J.R. Decker, A.J. Hull, J.M. Verboon, and S.M. Parkhurst. 2018. Into the breach: how cells cope with wounds. Open Biol. 8.

Nakamura, M., J.M. Verboon, T.E. Allen, M.T. Abreu-Blanco, R. Liu, A.N.M. Dominguez, J.J. Delrow, and S.M. Parkhurst. 2020. Autocrine insulin pathway signaling regulates actin dynamics in cell wound repair. PLoS Genet. 16:e1009186.

Nakamura, M., J.M. Verboon, and S.M. Parkhurst. 2017. Prepatterning by RhoGEFs governs Rho GTPase spatiotemporal dynamics during wound repair. J Cell Biol. 216:3959–3969.

Naraghi, M., and E. Neher. 1997. Linearized buffered Ca2+ diffusion in microdomains and its implications for calculation of [Ca2+] at the mouth of a calcium channel. J Neurosci. 17:6961–6973.

Ni, J.-Q., R. Zhou, B. Czech, L.-P. Liu, L. Holderbaum, D. Yang-Zhou, H.-S. Shim, R. Tao, D. Handler, P. Karpowicz, R. Binari, M. Booker, J. Brennecke, L.A. Perkins, G.J. Hannon, and N. Perrimon. 2011. A genome-scale shRNA resource for transgenic RNAi in Drosophila. Nature methods. 8:405–407.

Pervin, M.S., G. Itoh, M.S.U. Talukder, K. Fujimoto, Y.V. Morimoto, M. Tanaka, M. Ueda, and S. Yumura. 2018. A study of wound repair in Dictyostelium cells by using novel laserporation. Sci Rep. 8:7969.

Potez, S., M. Luginbuhl, K. Monastyrskaya, A. Hostettler, A. Draeger, and E.B. Babiychuk. 2011. Tailored protection against plasmalemmal injury by annexins with different Ca2+ sensitivities. J Biol Chem. 286:17982–17991.

Quattrocelli, M., J. Capote, J.C. Ohiri, J.L. Warner, A.H. Vo, J.U. Earley, M. Hadhazy, A.R. Demonbreun, M.J. Spencer, and E.M. McNally. 2017. Genetic modifiers of muscular dystrophy act on sarcolemmal resealing and recovery from injury. PLoS Genet. 13:e1007070.

Rescher, U., C. Ludwig, V. Konietzko, A. Kharitonenkov, and V. Gerke. 2008. Tyrosine phosphorylation of annexin A2 regulates Rho-mediated actin rearrangement and cell adhesion. Journal of cell science. 121:2177–2185.

Ritter, A.T., G. Shtengel, C.S. Xu, A. Weigel, D.P. Hoffman, M. Freeman, N. Iyer, N. Alivodej, D. Ackerman, I. Voskoboinik, J. Trapani, H.F. Hess, and I. Mellman. 2022. ESCRT-mediated membrane repair protects tumor-derived cells against T cell attack. Science. 376:377–382.

Rosales-Nieves, A.E., J.E. Johndrow, L.C. Keller, C.R. Magie, D.M. Pinto-Santini, and S.M. Parkhurst. 2006. Coordination of microtubule and microfilament dynamics by Drosophila Rho1, Spire and Cappuccino. Nat Cell Biol. 8:367–376.

Schindelin, J., I. Arganda-Carreras, E. Frise, V. Kaynig, M. Longair, T. Pietzsch, S. Preibisch, C. Rueden, S. Saalfeld, B. Schmid, J.Y. Tinevez, D.J. White, V. Hartenstein, K. Eliceiri, P. Tomancak, and A. Cardona. 2012. Fiji: an open-source platform for biological-image analysis. Nat Methods. 9:676–682.

Sønder, S.L., T.L. Boye, R. Tölle, J. Dengjel, K. Maeda, M. Jäättelä, A.C. Simonsen, J.K. Jaiswal, and J. Nylandsted. 2019. Annexin A7 is required for ESCRT III-mediated plasma membrane repair. Scientific Reports. 9:6726.

Sonnemann, K.J., and W.M. Bement. 2011. Wound Repair: Toward Understanding and Integration of Single-Cell and Multicellular Wound Responses. Annual Review of Cell and Developmental Biology. 27:237–263.

Spradling, A.C. 1986. P element-mediated transformation. In In Drosophila: A practical approach. D.B. Roberts, editor. IRL Press, Oxford. 175–197.

Talukder, M.S.U., M.S. Pervin, M.I.O. Tanvir, K. Fujimoto, M. Tanaka, G. Itoh, and S. Yumura. 2020. Ca(2+)-Calmodulin Dependent Wound Repair in Dictyostelium Cell Membrane. Cells. 9.

Terasaki, M., K. Miyake, and P.L. McNeil. 1997. Large plasma membrane disruptions are rapidly resealed by Ca2+-dependent vesicle-vesicle fusion events. J Cell Biol. 139:63–74.

Tsurumi, F., S. Baba, D. Yoshinaga, K. Umeda, T. Hirata, J. Takita, and T. Heike. 2019. The intracellular Ca2+ concentration is elevated in cardiomyocytes differentiated from hiPSCs derived from a Duchenne muscular dystrophy patient. PLoS One. 14:e0213768.

Vaughan, E.M., J.S. You, H.Y. Elsie Yu, a. Lasek, N. Vitale, T.a. Hornberger, and W.M. Bement. 2014. Lipid domain-dependent regulation of single-cell wound repair. Molecular Biology of the Cell. 25:1867–1876.

Vicic, N., X. Guo, D. Chan, J.G. Flanagan, I.A. Sigal, and J.M. Sivak. 2022. Evidence of an Annexin A4 mediated plasma membrane repair response to biomechanical strain associated with glaucoma pathogenesis. J Cell Physiol. 237:3687–3702.

Wirtz, D., K. Konstantopoulos, and P.C. Searson. 2011. The physics of cancer: the role of physical interactions and mechanical forces in metastasis. Nat Rev Cancer. 11:512–522.

Xu, M., A. Seas, M. Kiyani, K.S.Y. Ji, and H.N. Bell. 2018. A temporal examination of calcium signaling in cancer-from tumorigenesis, to immune evasion, and metastasis. Cell & Bioscience. 8:1–9.

Zhang, F., Z. Wang, M. Lu, Y. Yonekubo, X. Liang, Y. Zhang, P. Wu, Y. Zhou, S. Grinstein, J.F. Hancock, and G. Du. 2014. Temporal Production of the Signaling Lipid Phosphatidic Acid by Phospholipase D2 Determines the Output of Extracellular Signal-Regulated Kinase Signaling in Cancer Cells. In Mol Cell Biol. Vol. 34. 84–95.

Zhou, Y., J. Shi, J. Cui, and C.X. Deng. 2008. Effects of extracellular calcium on cell membrane resealing in sonoporation. J Control Release. 126:34–43.

## REFERENCES

Schindelin, J., I. Arganda-Carreras, E. Frise, V. Kaynig, M. Longair, T. Pietzsch, S. Preibisch, C. Rueden, S. Saalfeld, B. Schmid, J.-Y. Tinevez, D.J. White, V. Hartenstein, K. Eliceiri, P. Tomancak, and A. Cardona. 2012. Fiji: an open-source platform for biological-image analysis. Nat Meth. 9:676–682.

